# Optogenetic manipulation of YAP cellular localisation and function

**DOI:** 10.1101/2021.03.19.436118

**Authors:** P. J. Y. Toh, J. K. H. Lai, A. Hermann, O. Destaing, M. P. Sheetz, M. Sudol, T. E. Saunders

## Abstract

YAP, an effector of the Hippo signalling pathway, promotes organ growth and regeneration. Prolonged YAP activation results in uncontrolled proliferation and cancer. Therefore, exogenous regulation of YAP activity has potential translational applications. We present a versatile optogenetic construct (optoYAP) for manipulating YAP localisation, and consequently its activity and function. We attached a LOV2 domain that photocages a nuclear localisation signal (NLS) to the N-terminus of YAP. In 488 nm light, the LOV2 domain unfolds, exposing the NLS, which shuttles optoYAP into the nucleus. Nuclear import of optoYAP is reversible and tuneable by light intensity. In cell culture, activated optoYAP promotes YAP target gene expression, cell proliferation, and anchorage-independent growth. Similarly, we can utilise optoYAP in zebrafish embryos to modulate target genes. OptoYAP is functional in both cell culture and *in vivo*, providing a powerful tool to address basic research questions and therapeutic applications in regeneration and disease.

## Introduction

The Hippo signalling pathway regulates organ size control and cell fate during development and regeneration.^1^ Components of this pathway were identified in a *Drosophila melanogaster* genetic screen^2,3^ and are evolutionary conserved.^4,5^ The core kinases in the Hippo pathway are MST1/2 and LATS1/2. MST1/2 phosphorylates and activates LATS1/2^6^ in the presence of SAV1^7^ and MOB1A/B.^8^ Subsequently, activated LATS1/2 phosphorylates YAP/TAZ^3^ which results in their cytoplasmic sequestration from the nucleus by 14-3-3 proteins^9,10^ or their degradation.^11^

Cellular responses to Hippo signalling include proliferation, migration, changes in cellular cytoskeleton and morphology, as well as cell survival.^12^ Genetic loss of the Hippo kinases or hyperactivation of YAP/TAZ (Yorkie in *D. melanogaster*), has been shown to promote tissue regeneration,^13–15^ but at the risk of organ overgrowth.^4,16,17^ Furthermore, YAP/TAZ are also key drivers in tumour development.^18^ Therefore, to harness the therapeutic benefits of YAP/TAZ without suffering from their tumorigenic potential, developing a tool to control the activation of YAP is desirable.

Optogenetics is a powerful method for controlling the spatiotemporal localisation and activation of specific biological processes. It has been used extensively to activate neural responses.^19^ Optogenetic strategies have also been deployed to modulate and probe cell signalling pathways^20^ where high spatiotemporal control is desirable, such as Ras/Erk,^21^ Akt,^22^ Notch,^23^ Bicoid,^24^ Src^25^ and p53 signalling pathways.^26^ In light of the variety of optogenetic tools and capabilities, optogenetic approaches potentially enable precise regulation of the YAP/TAZ nuclear-cytoplasmic localisation, which can modulate their co-transcriptional activity.

To this end, we utilised the LOV2-Jα interacting domain to photocage a NLS within the Jα helix domain.^27^ Fusing this light inducible domain to the N-terminus of YAP, henceforth referred to as optoYAP, thus enables manipulation of YAP cellular localisation with light, which we report here. We show that optoYAP is imported into the nucleus after a 488 nm light activation protocol in a range of cell lines and in the zebrafish embryo. We find that the optoYAP nuclear import dynamics are tuneable by activation light intensity. Activated optoYAP elicits cellular responses including the expression of downstream target genes and cell proliferation. Strikingly, we find that optoYAP can promote anchorage-independent growth, showing that optoYAP activation is sufficient to override the mechano-inhibitory signals in cells seeded on soft growing substrates. Such specific control opens new opportunities to explore key questions concerning dysregulation of YAP localisation and its effects on disease progression. Further, it enables the potential for the manipulation of YAP/TAZ activity in therapeutic applications.

## Results

### Optogenetic YAP construct

To tightly control YAP localisation both temporally and spatially, we fused an optogenetic construct to the N-terminal end of hYAP1.^27^ This construct consists of the LOV2 domain of *Avena sativa* phototropin 1 with mCherry for visualisation (Fig. 1A). The LOV2 domain is connected to hYAP1 via a Jα helix and NLS sequence. A nuclear export signal (NES) derived from truncated cAMP-dependent protein kinase (PKI) facilitates nuclear export and decreases background nuclear localisation in the dark state.^27^ We refer to this construct as optoYAP (Fig. 1A).

**Figure 1.**
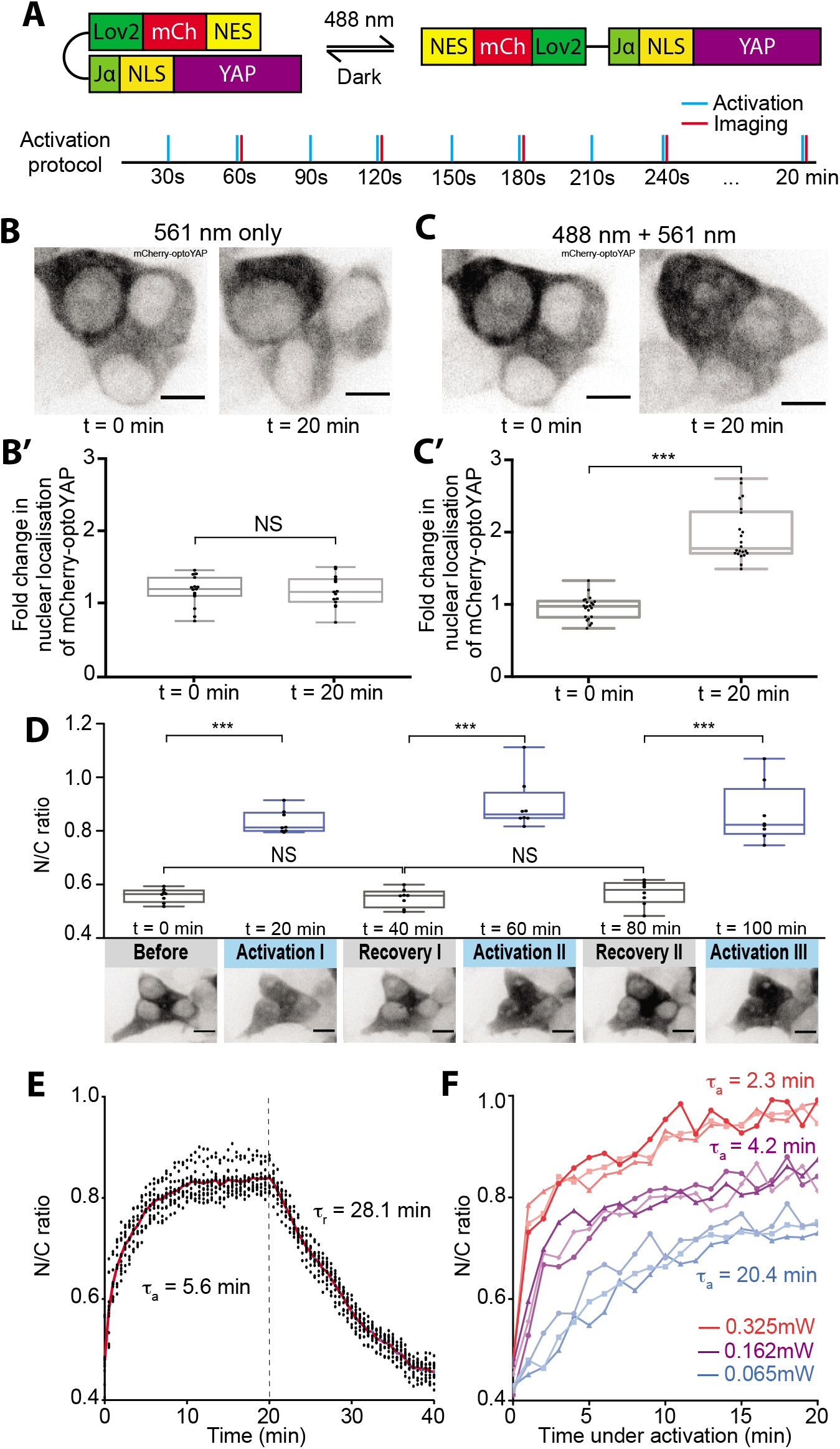
Characterisation of optoYAP in cell culture. **(A)** Schematic of optoYAP construct. OptoYAP is folded in the dark due to the interaction between LOV2 domain and Jα helix. Under 488 nm light, nuclear localisation signal (NLS) is exposed and optoYAP is transported into the nucleus. Activation protocol was performed with 1 s blue light pulse every 30 s (blue bars), and cells were imaged every minute (red bars) for 20 minutes. **(B)** Representative images of mCherry-optoYAP in HEK293T cells imaged with only 561 nm laser following the protocol in (A). **(B’)** Fold-change in nuclear localisation of mCherry-optoYAP (n=22 cells from 2 independent experiments). **(C)** Representative images of the same cells in (B) exposed to both 488 and 561 nm following the protocol shown in (A). **(C’)** Fold-change in nuclear localisation of the mCherry-optoYAP (n=22 cells from 2 independent experiments). **(D)** HEK293T cells transfected with optoYAP were subjected to three cycles of activation protocol and recovery in the dark. (Top) The ratio of mCherry-optoYAP signal in the nucleus to the cytoplasmic mCherry-optoYAP signal (n=8 cells from 2 independent experiments). (Bottom) Representative images of the mCherry-optoYAP signal from the same HEK293T cells at each cycle. **(E)** Activation time constant (ta) and recovery time constant (tr) (see Methods and Fig. S1B-C) of optoYAP in HEK293T cells. Vertical dashed line represents time when 488 nm stimulation ceased. Red line indicates average nuclear/cytoplasmic ratio (n=12 cells). **(F)** HEK293T cells transfected with optoYAP subjected to 0.065, 0.163 and 0.325 mW 488 nm laser power at the focal plane. n=3 cells per laser power. NS: Not significant, \*\*\**P* < 10^−3^. Box plots represent median and 25^th^ to 75^th^ percentiles. Bars show minimum and maximum points.

In the dark, the LOV2 domain photocages the NLS^27^ to prevent importation of optoYAP into the nucleus (Fig. 1A). Under blue light illumination, the Jα helix domain unfolds and is released from the LOV2 domain to expose the NLS and allows transport of optoYAP into the nucleus.

### Characterisation of optogenetic YAP in mammalian cells

HEK293T cells were transiently transfected with optoYAP and subjected to 488 nm pulsatile illumination of 1 s every 30 s, similar to optogenetic activation protocols described previously^27^ (Fig. 1A). Localisation of optoYAP is then visualised by the attached mCherry fluorophore. Whereas optoYAP is localised to the cytoplasm without activation (Fig. 1B-B’), its accumulation in the nucleus doubles after 20 minutes of activation (Fig. 1C-C’ and Supplementary Video 1).

We tested whether optoYAP shuttles in and out of the nucleus by recurring activation cycles. We subjected HEK293T cells to three rounds of activation protocol (20 minutes) and recovery in the dark (20 minutes). OptoYAP accumulates in the nucleus to the same level when activated and retreats to its baseline level after each activation cycle (Fig. 1D and Fig. S1A). This property shows that optoYAP activation is reversible and its light-sensitive property is not attenuated over activation cycles.

To explore the nuclear import and export dynamics of optoYAP, we performed time-lapse imaging of optoYAP in HEK293T cells. Cells were exposed to the above activation protocol for 20 minutes and then the cells were left in the dark for a further 20 minutes, while optoYAP localisation was tracked every 30 seconds. The nuclear accumulation saturated quickly, with an activation time constant (ta) of 5.6±0.4 minutes (Fig. 1E, Fig. S1B and Methods). The recovery time constant (tr) was 28.1±2.4 minutes (Fig. 1E and Fig. S1C).

We next tested the light sensitivity of optoYAP in HEK293T cells. We performed a series of light activations using varying 488 nm laser power, 0.065, 0.162, and 0.325 mW, measured at the focal plane. Both ta (Fig. S1D) and extent of nuclear accumulation of optoYAP positively responded to increasing laser powers (Fig. 1F).

Phosphorylation of YAP at serine 127 residue has been shown to promote its cytoplasmic localisation.^28^ We investigated the phosphorylation status of optoYAP when it is nuclear localised after light activation. Using CRISPR-mediated *YAP* knockout MKN28 cells, we transfected optoYAP into these cells and probed for pYAP (S127) using Western blot (Fig. S1E). Phosphorylated optoYAP was clearly present in the nucleus, suggesting that the NLS in our optoYAP construct can import phosphorylated optoYAP marked for cytoplasmic sequestration.

### Functionality of nuclear-localised optoYAP

We next investigated the functionality of optoYAP. As YAP is a transcriptional co-regulator, it binds to TEAD transcription factors in the nucleus to initiate transcription of downstream genes.^29^ HEK293T cells transiently transfected with optoYAP were subjected to pulsed light activation over 48 h followed by qPCR assays of YAP target genes: *ANKRD1, CTGF* and *CYR61*. The expression levels of all three genes were significantly upregulated in cells with activated optoYAP construct as compared to control cells without light activation (Fig. 2).

**Figure 2.**
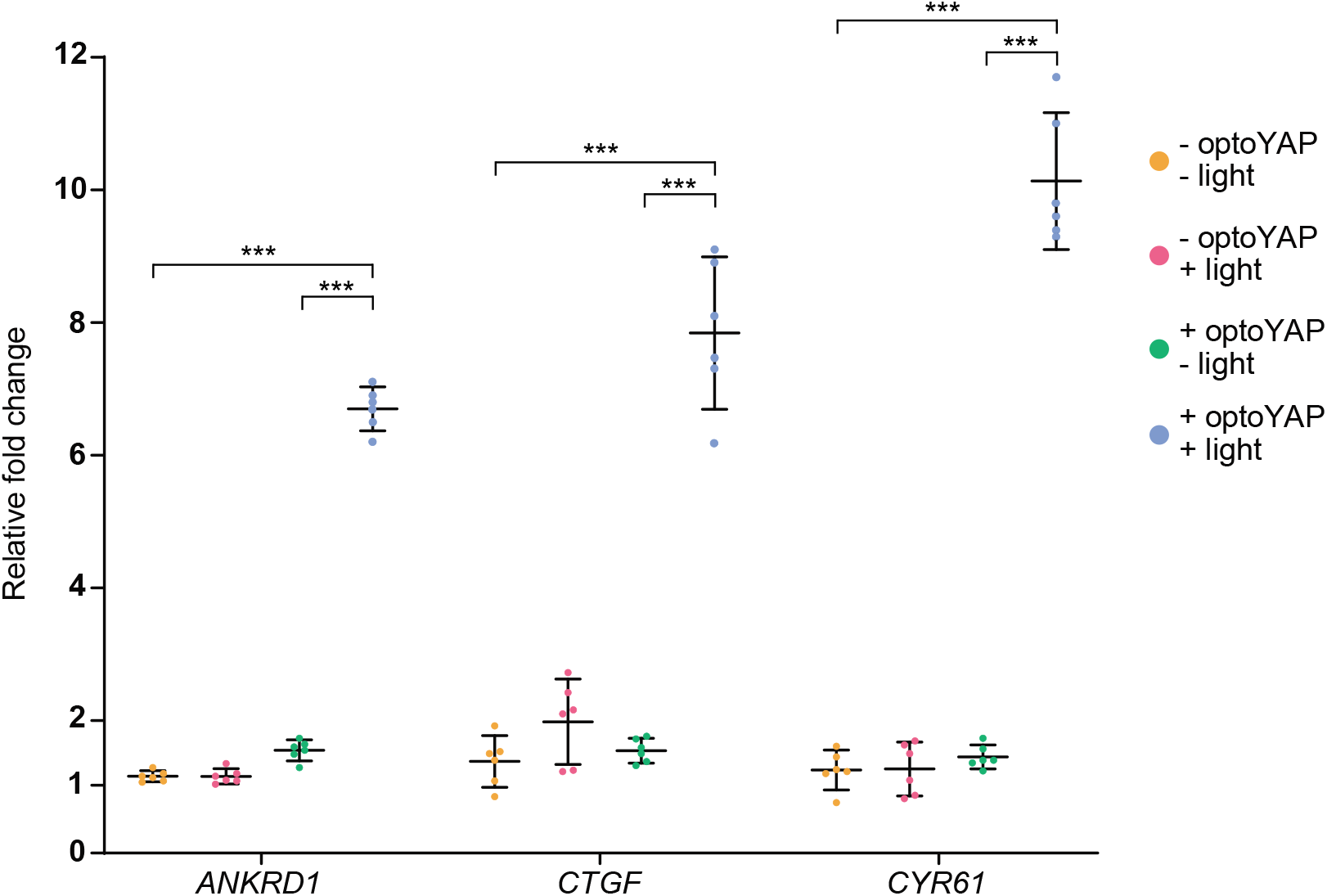
qPCR data of downstream YAP target genes. Expression levels of *ANKRD1, CTGF*, and *CYR61* transcripts in HEK293T cells transfected with optoYAP after 48 h of activation protocol. Gene expression level was normalized to the housekeeping gene, *EIF1B*. Horizontal bars represent mean and 95% confidence interval from 6 biological replicates across 2 independent experiments for each condition, \*\*\**P* < 10^−3^.

Given that activated optoYAP can activate downstream targets, we postulated that activated optoYAP could also promote cells to proliferate. We assayed cell proliferation of two cell lines, HEK293T and HFF, by using a DNA-binding fluorescent dye to measure their total DNA. Cells were transfected with optoYAP and subjected to pulsatile light activation over one week.

Both transformed (HEK293T, Fig. 3A) and non-transformed (HFF, Fig. 3B) cell lines show significant increase in total DNA with activated optoYAP as compared to untransfected and unactivated controls.

**Figure 3.**
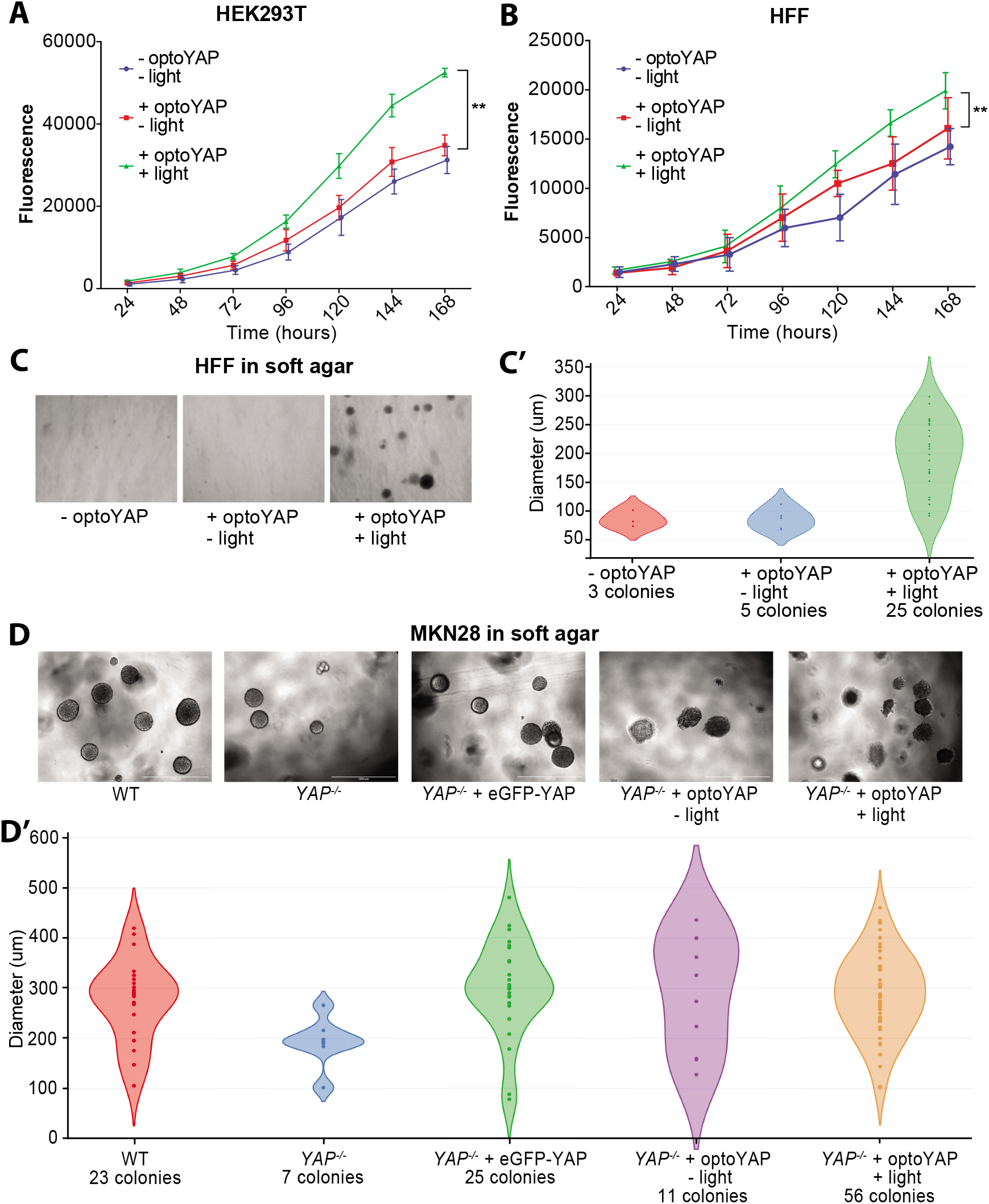
Functional assays of optoYAP in tissue culture cells. **(A-B)** Cell proliferation assay. HEK293T **(A)** and HFF **(B)** cells were transfected with optoYAP and subjected to activation protocol as described in Fig. 1A or kept in the dark for one week. Fluorescence measures amount of DNA present in each well (Methods). Error bars are s.d., n=3 independent experiments for each condition, \*\**P* < 10^−2^. **(C-D)** Soft agar colony formation assay. HFF (**C**) and MKN28 (**D**) cells grown on soft agar for one week under different light conditions as described in Fig. 1A. (**C’-D’**) Diameter of individual colonies formed under different light conditions and genetic backgrounds. Colonies were counted and measured from 3 biological replicates.

### optoYAP can induce anchorage independent growth

Mechanical parameters of the microenvironment, such as matrix rigidity, are crucial for normal cell growth.^30^ In particular, epithelial cells typically cannot survive and proliferate on soft substrates. However, transformed cells are able to bypass rigidity sensing and grow on soft substrates.^31^ We hypothesised that activated optoYAP may drive proliferation of cells even on soft substrates. Using a non-transformed cell line, HFF, we cultured untransfected and optoYAP-transfected cells with or without light activation on soft agar for seven days. As expected, few untransfected cells survived, with little visible colony formation (Fig. 3C). However, optoYAP-transfected cells that were subjected to light activation formed more colonies (Fig. 3C and Fig. S2A) with larger size (Fig. 3C’) compared to those without light activation. The increase in number and size of colonies formed indicates that activated optoYAP can induce transformed growth, overriding the ability of normal cells to sense their microenvironment and suppress growth in soft substrates.

Similarly, we performed the soft agar assay on a transformed cell line MKN28. Using *YAP^-/-^* MKN28 cells,^32^ we rescued these mutant cells with either eGFP-YAP or optoYAP. We seeded four variants of MKN28 cells – wildtype (WT), *YAP^-/-^*, *YAP^-/-^* + eGFP-YAP, and *YAP^-/-^* + optoYAP – on soft agar for seven days. WT MKN28 cells, unlike HFF cells, are transformed cells and are able to form colonies on soft agar, which is attenuated by the loss of *YAP* (Fig. 3D-3D’ and Fig. S2B). While overexpression of eGFP-YAP rescues *YAP* null MKN28 cells to form colonies at WT levels, activated optoYAP exaggerates colony formation of MKN28 cells on soft agar (Fig. 3D and Fig. S2B). Protein expression levels of both eGFP-YAP and optoYAP transfected in *YAP* null cells were similar (Fig. S2C). However, optoYAP was localised in the nucleus during the activation protocol, therefore likely inducing cell proliferation and increasing the number of colonies formed (Fig. 3D’ and Fig. S2B). The colony size did not vary between the different MKN28 cell lines, with the exception of *YAP* null cells, but the number of colonies formed with activated optoYAP is more than double of the WT or eGFP-YAP rescue (Fig. S2B). These data show that exogenous and sustained activation of optoYAP in MKN28 cells triggers greater levels of cell growth on soft agar than that of WT or rescued *YAP* mutant cells.

### optoYAP in zebrafish embryos

Building on the functional results in mammalian cell culture, we tested optoYAP in an *in vivo* system: the developing zebrafish embryo. We replaced hYAP by fYap; we refer to this construct as optofYap. Shield stage zebrafish embryos expressing *optofYap* mRNA were imaged at the animal pole (Fig. 4A). optofYap is distributed uniformly in the entire cell (though we observed mosaic levels of expression between cells), prior to activation (Fig. 4B). Using a 488 nm pulsatile light activation protocol as per Fig. 1A, we see that optofYap can shuttle into the nuclei of both the enveloping layer (EVL) and deep cells (Fig. 4B and Supplementary Video 2). We observe a two-fold change in nuclear localisation after light activation (Fig. 4B’). Interestingly, optofYap appears to enter and leave the nucleus much faster in the zebrafish embryos than optoYAP does in cultured cells. The activation time constant was calculated to be 2.9±0.8 minutes and recovery time constant at 4.8±1.6 minutes (Fig. 4C and Fig. S3A-B).

**Figure 4.**
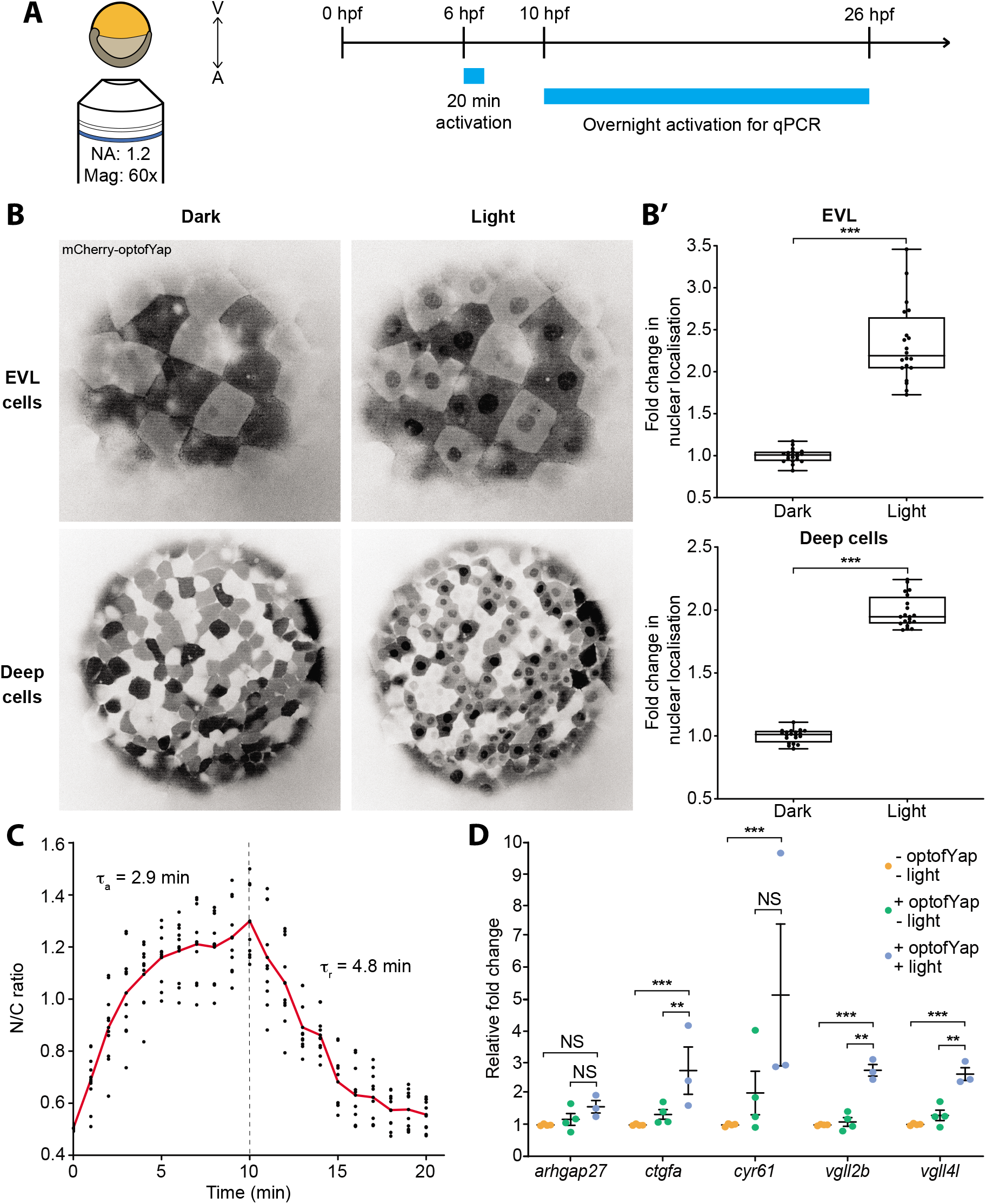
Validation of optofYap in zebrafish. **(A)** Imaging and light activation protocol in zebrafish. Shield-stage embryos were subjected to activation protocol as described in Fig. 1A and imaged at the animal pole. **(B)** Example images of embryos expressing optofYap (mCherry-tagged) in the EVL and deep cells kept in the dark or subjected to the activation protocol at 6 hpf. **(B’)** Fold-change in nuclear localisation of mCherry-optofYap (n=20 cells from 3 independent experiments). Box plots represent median and 25^th^ to 75^th^ percentiles. Whiskers show minimum and maximum points. **(C)** The N/C ratio measured using mCherry signal in zebrafish embryos injected with optofYap tracked over 10 min of activation protocol and 10 min recovery in the dark (see also Fig. S3A-B). (n=10 cells from 2 independent experiments, red line=average). Vertical dashed line represents time when 488 nm stimulation ceased. **(D)** qPCR of Yap target genes. Embryos were kept in the dark until 10 hpf and then subjected to pulsed light activation for 16 h. Gene expression level was normalized to the housekeeping gene, *rpl13*. Horizontal bars represent mean and 95% confidence interval from 4 biological replicates (NS: Not significant, \*\**P* < 10^−2^, \*\*\**P* < 10^−3^).

Similar to our tissue culture studies, we tested if optofYap can induce the expression of specific Yap target genes in the embryo after activation.^33^ qPCR was performed on embryos subjected to overnight activation protocol for 14 hours from the tailbud stage (10 hours post fertilisation (hpf)). Expression of *arhgap27* did not change in all conditions, while *ctgfa, vgll2b and vgll4b* were upregulated after activation. Even though *cyr61* was upregulated by optofYap, contribution by the activation protocol is unclear (Fig. 4D).

## Discussion

We report the development of a functional optogenetic YAP, optoYAP, by exploiting the light-responsive LOV2-Jα domain to photocage NLS. We characterised optoYAP function in human cells and zebrafish embryos, showing a consistent increase in YAP nuclear localisation upon blue light illumination within minutes. This rapid manipulation of YAP localisation elicits downstream responses, including the upregulation of target genes, increased cell proliferation in cell culture, and anchorage-independent growth. Long term activation of optoYAP does not appear to have detrimental effects in cell culture. Each activation cycle is independent (Fig. 1D), and cells survive after being subjected to one week of light activation. This adds to the advantage of optogenetic YAP over constitutively active YAP or drug-induced YAP expression.

A similar optogenetic YAP based on the LOV2-TRAP tool has recently been described.^34^ However, the construct does not show a functional YAP response upon activation. There, LOV2 is tethered on the outer surface of the mitochondria and YAP is tagged with a small peptide Zdk which interacts with the LOV2 domain.^34^ This system requires two-plasmids to be introduced into the tissue of interest. Our optogenetic YAP construct described here requires only a single component, the LOV2-Ja interacting domain, thus simplifying its deployment and presents an advantage for future applications. Activation of the photosensitive domain can be achieved within visible light range at 488 nm and recovery done in the dark. These features add to the ease of use over other optogenetic approaches that require multiple components or activation in the infrared spectrum.^35^ Further, to the best of our knowledge, this is the first report of a functional optogenetic YAP construct.

We observed two differences in our optogenetic construct between tissue culture cells and zebrafish embryos. Firstly, without activation, optofYap is uniformly distributed in zebrafish cells (Fig. 4B), in contrast to optoYAP which is localized to the cytoplasm of human cell lines (Fig. 1C). Secondly, the activation time constant is faster in zebrafish (2.9 min, Fig. 4C) than in cultured cells (5.6 min, Fig. 1E). The NLS and NES present in this optogenetic construct were optimized from human c-Myc and PKI, respectively.^27^ These two peptide sequences differ from their respective *Danio rerio* orthologue (Fig. S3C-D). This sequence divergence suggests that the nuclear import/export machineries in the zebrafish may be more sensitive to modified c-Myc NLS and could explain the differing dynamics of optoYAP between human and zebrafish cells. The nuclear import and export rates also differ in cell culture (Fig. 1E), which suggests that here the rate of nuclear import driven by the unmasked NLS is greater than the rate of nuclear export driven by the NES.

We found that a transformed cell line, MKN28, depends on YAP for anchorage-independent growth, as the removal of *YAP* results in fewer and smaller colonies in the soft agar assay. Moreover, forced overexpression of eGFP-YAP in these mutant cells reinstated colony growth to similar to WT cell levels. On the other hand, light activation of optoYAP further amplified cell proliferation by increasing the number of colonies formed. One possibility is that eGFP-YAP is limited by upstream signals including Hippo kinases, and that our optoYAP appears to override such limitations as seen in our Western blot data (Fig. S1E). This shows that optoYAP might be able to bypass canonical Hippo signalling regulation.

There is a potential for induced activation of YAP to promote regeneration of injured organs.^36^ However, sustained activation can lead to uncontrolled proliferation.^37^ YAP has been shown to be required for liver regeneration after partial hepatectomy, where regenerating hepatocytes had increased YAP activity that returned to original levels once liver size was restored.^38^ On the other hand, in an organ with low regenerative capacity like the heart, overexpression of YAP is sufficient to promote myocardial proliferation and cardiac regeneration.^15,39^ Although constitutively-active YAP in cardiomyocytes increases ventricular wall thickness and ejection fraction, its chronic activity is fatal.^40^ Therefore, heart regeneration represents an opportunity for the application of optoYAP to limit the activity of YAP for cardiomyocyte renewal.

Overall, given the importance of YAP with respect to development and regeneration, and the ease of the optogenetic tool presented here, optoYAP has the potential to become a powerful tool for studying Hippo-YAP signalling in both research and clinical applications.

## Acknowledgements

This work was supported by a Singapore Ministry of Education Tier 3 grant (MOE2016-T3-1-002). We thank Richard De Mets for his help on data analysis. We thank members of the Saunders, Sudol, and Sheetz labs for feedback on the project.

## Author Contributions

P.J.Y.T., M.S. and M.P.S. conceived the study. P.J.Y.T., J.K.H.L., M.S., M.P.S., T.E.S. designed and planned the experiments. P.J.Y.T. performed the cell culture and zebrafish experiments. J.K.H.L. assisted with the experiments, particularly in the zebrafish work. A.H. generated the initial optoYAP clone with the help of O.D.. P.J.Y.T. analyzed the data with assistance from J.K.H.L. and T.E.S.. P.J.Y.T., J.K.H.L. and T.E.S. wrote the first draft of the manuscript with all authors approving the final manuscript.

---

**Figure S1.**
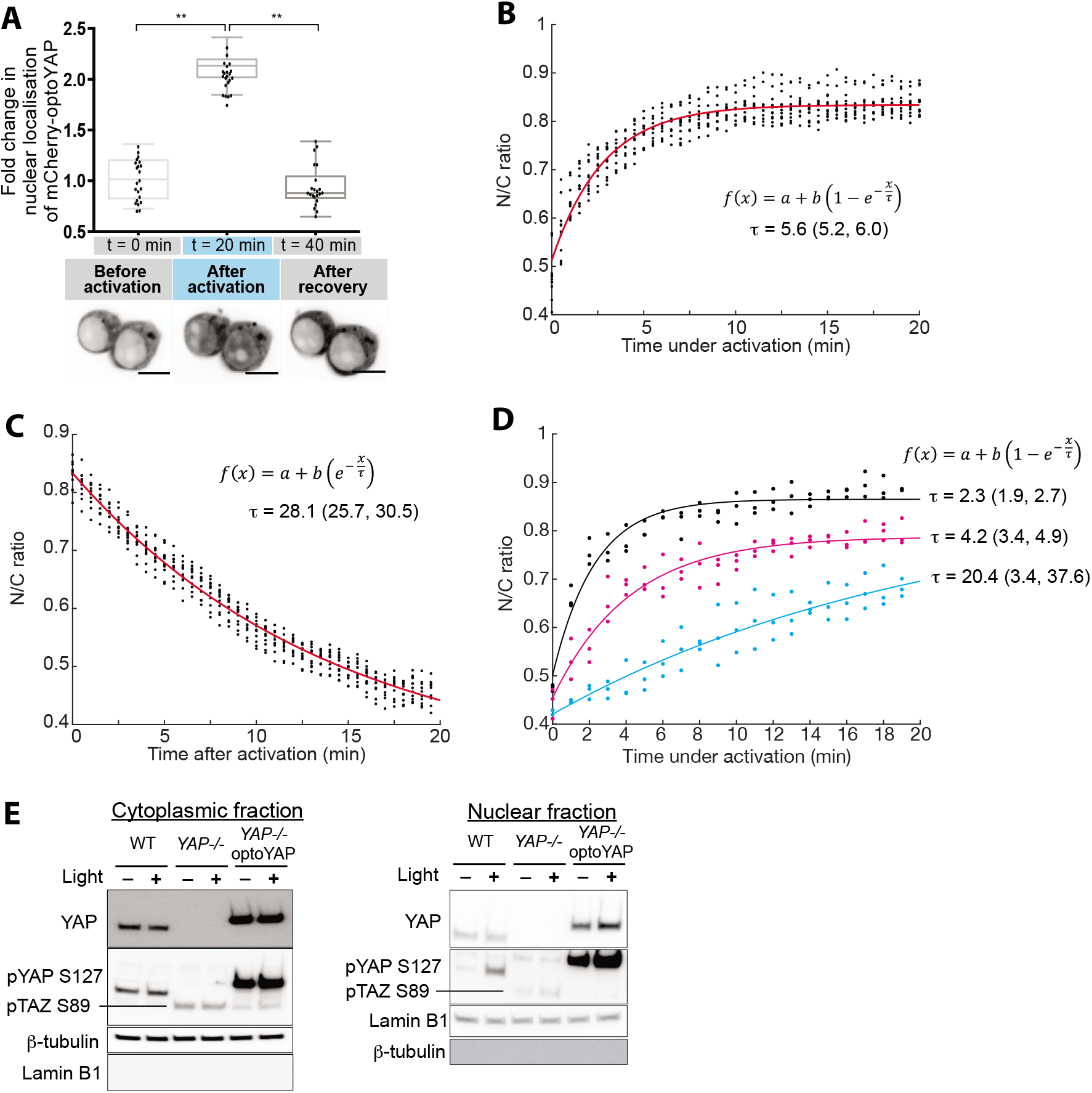
Characterisation of optoYAP in tissue culture cells. **(A)** HEK293T transfected with optoYAP were subjected to activation protocol in Fig. 1A followed by recovery in the dark for 20 min. Fold-change in nuclear localisation of mCherry-optoYAP (n=22 cells from 2 independent experiments). Box plots represent median and 25^th^ to 75^th^ percentiles. Bars show minimum and maximum points, \*\**P* < 10^−2^. **(B-C)** Curve fitting for Fig. 1E. Red line represents the exponential curve fitted to the data. Numbers in brackets represent the 95% confidence interval of τ. **(D)** Curve fitting for Fig. 1F for three different laser powers. Red line represents the exponential curve fitted to the data. Numbers in brackets represent the 95% confidence interval of τ. **(E)** Western blots of MKN28 cells. MKN28 WT, *YAP^-/-^*, and *YAP^-/-^* cells transfected with optoYAP were subjected to pulsed light activation for 48 h. Whole cell lysate from the three cell lines were separated into nuclear and cytoplasmic fractions, then probed for YAP, pYAP (S127), lamin B1 and β–tubulin.

**Figure S2.**
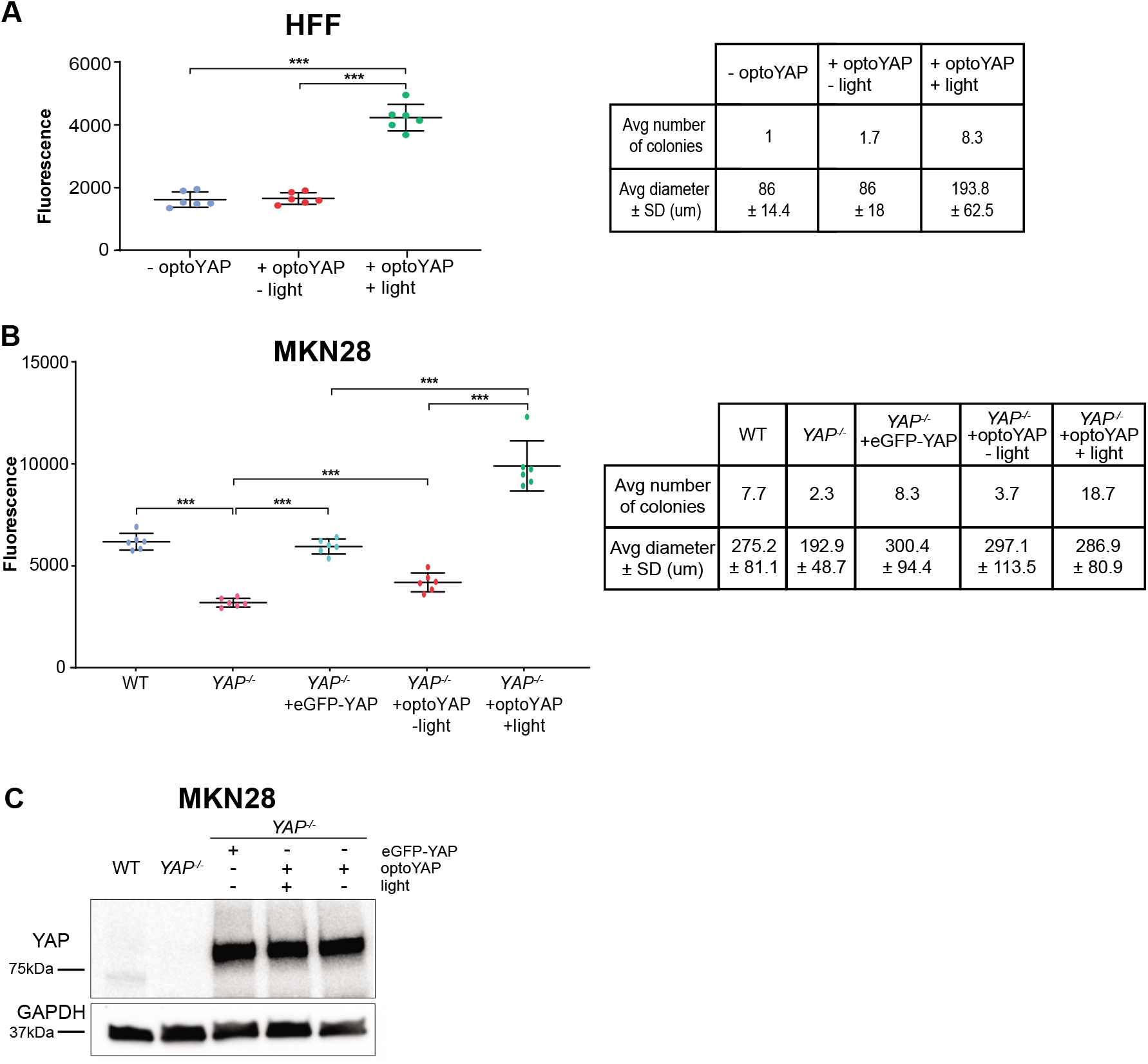
Colony formation assay. (**A-B)** Quantification of DNA-binding fluorescent dye in HFF **(A)** and MKN28 **(B)** cells grown on soft agar. The average number and diameter of colonies formed (representative images in Fig. 3C-D) are shown in the table on the right. Error bars are s.d., n=6 biological replicates from 2 independent experiments for each condition, \*\*\**P* < 10^−3^. **(C)** Western blots of different MKN28 cell lines. eGFP-YAP and optoYAP lines are transfected into the *YAP^-/-^* background.

**Figure S3.**
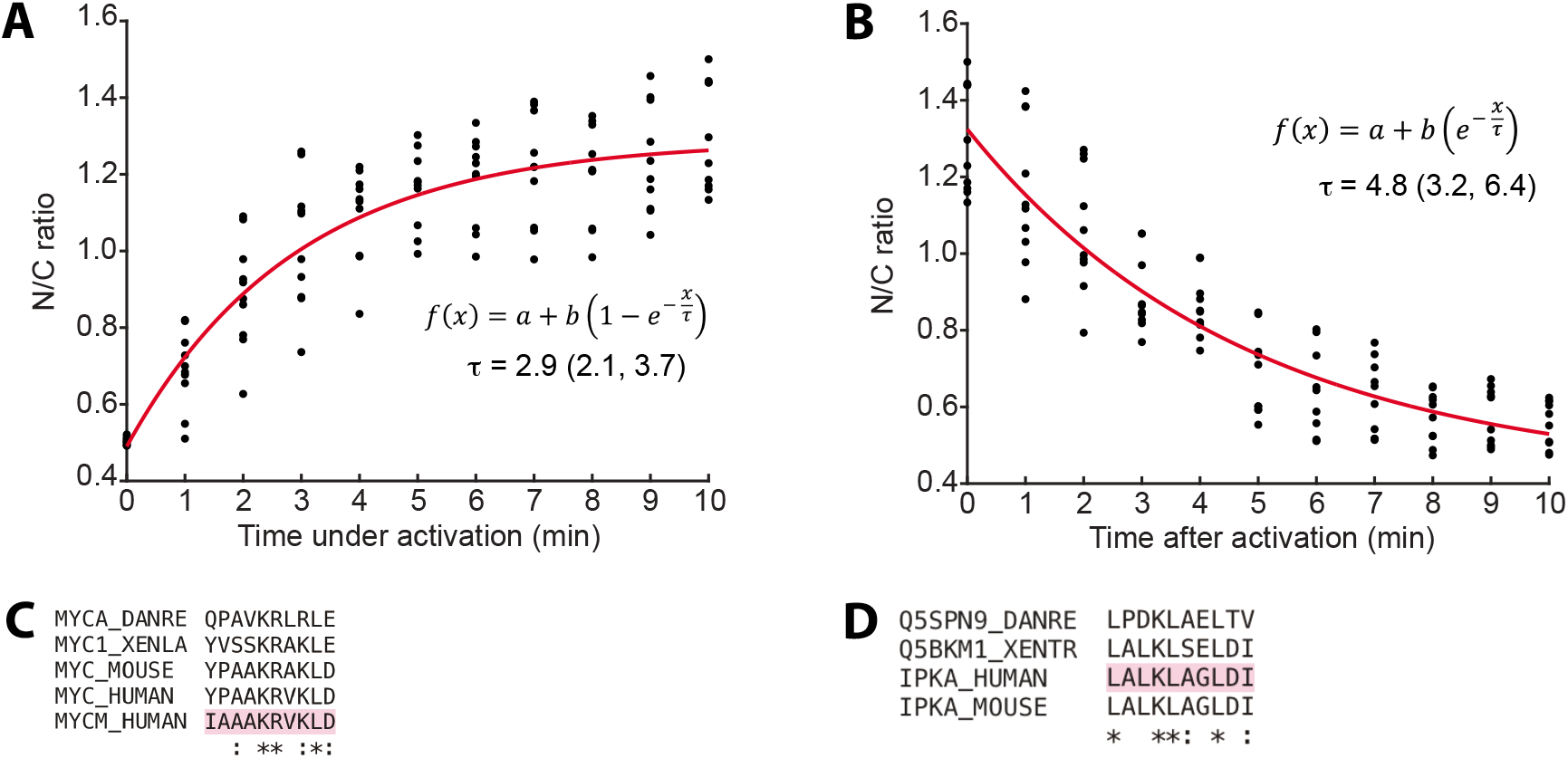
Characterisation of optofYAP in zebrafish embryos. **(A-B)** Curve fitting for Fig. 4C. Red line represents the exponential curve fitted to the data. Numbers in brackets represent the 95% confidence interval of τ. **(C-D)** Multiple sequence alignments for the NLS of c-Myc **(C)** and NES of PKI **(D)** between human, mouse, frog, zebrafish and the optogenetic construct backbone (highlighted in pink).

**Supplementary Video 1.** HEK293T cells transfected with optoYAP under 561 nm only (left) or 488 and 561 nm (right) activation.

**Supplementary Video 2.** Z-stack of zebrafish embryo expressing mCherry-optofYap (magenta) and eGFP-Yap (green) before (left) and after (right) activation protocol.

## Methods

### Plasmid construction

Optogenetic plasmid (pDN34, previously reported in ^27^) containing the LOV2-Jα domain was provided by Olivier Destaing. Human YAP1-1δ was PCR amplified from pcDNA3.1 hYAP1-1δ with primers AH05 and AH06. Phusion High Fidelity DNA polymerase (ThermoScientific) was used for PCR amplification. PCR product and pDN34 were cleaved with KpnI and PmeI FastDigest restriction enzymes (ThermoScientific) for cloning.

### Mammalian cell culture and transfection

HEK293T and HFF cells were kept in phenol red-free Dulbecco’s modified Eagle medium (DMEM) (Gibco) supplemented with 10% (V/V) heat inactivated fetal bovine serum (FBS) (Hyclone) and 1% (V/V) penicillin/streptomycin (Nacalai Tesque). MKN28 cells were kept in RPMI 1640 supplemented with 10% (V/V) heat inactivated FBS (Hyclone) and 1% (V/V) penicillin/streptomycin (Nacalai Tesque). All cells were maintained in 37°C incubator with 5% CO2.

HEK293T and HFF cells were transfected with Lipofectamine 2000 (Invitrogen) according to the manufacturer’s instructions. MKN28 cells were transfected using K2 transfection system (Biontex).

### Zebrafish strains

All zebrafish strains were maintained according to standard fish husbandry procedures. The AB wild-type strain was used in this study. All experiments with zebrafish were approved by the A*STAR Biological Resource Centre according to the Singapore National Advisory Committee on Laboratory Animal Research.

### Zebrafish microinjection

*optofYap* was cloned into a pCS2 vector backbone by restriction enzyme cloning with EcoRI and PspXI (ThermoScientific) from optoYAP. The plasmid was linearised using NotI (ThermoScientific) and capped mRNA was synthesised using mMessage Machine SP6 kit (Ambion). Embryos from AB zebrafish were collected at one-cell stage and injected with 10 pg *optofYAP* mRNA.

### Fluorescence microscopy

Cells were visualised under the microscope about 24 h post transfection. Live imaging was performed on cells seeded on Iwaki glass-bottom dish and imaged on Perkin Elmer spinning disk with a LUCPlanFLN 40x/0.6 NA air objective. Injected zebrafish embryos were collected about 6h post fertilisation at shield stage, dechorionated and embedded in 1% low melting agarose dissolved in egg media. Embryos were mounted on a MatTek dish and imaged on Nikon Ti-E with Yokogawa W1 spinning disk with a PLAN APO VC 60X/1.40 NA oil immersion objective.

The Nuclear/cytoplasmic ratio was measured using Fiji by drawing a region of interest (ROI) around the cell as well as its nucleus. A macro plugin was used to mask the nucleus from the cytoplasm to calculate the mean intensity in the nucleus and cytoplasm based on the ROI drawn. Nucleus was demarcated by region without mCherry signal in cells at t = 0 min and region with highest level of mCherry in zebrafish embryos at t = 20 min. Laser power at the focal point was measured using a power meter (Thorlabs).

### Light activation

Pulsed light activation is as follows: cells are illuminated with a 1 s pulse of 488 nm laser light, followed by a 30 s dark phase. This pulsation is continued over 20 min on the Perkin Elmer or Nikon W1. For long term cell proliferation (7 d), soft agar assay (7 d) and qPCR experiments (48 h for tissue culture cells, 16 h for zebrafish embryos), cells were kept in cell culture incubators fitted with a LED light strip that pulsed at the same frequency.

### RT-qPCR

Total RNA was isolated from mammalian cell culture after 48 h of pulsatile light activation using RNeasy Mini Kit (Qiagen) according to the manufacturer’s instructions. Complementary DNA (cDNA) was synthesised from isolated RNA using SuperScript IV Reverse Transcriptase (ThermoScientific). Zebrafish embryos injected with optofYAP at the one cell stage were subjected to pulsatile light activation and collected after 48 h.

For zebrafish, 5 embryos were pooled into one biological replicate and a total of 4 biological replicates were obtained for each sample. RNA was extracted using TRIzol (Ambion) and purified with Direct-zol RNA kit (Zymogen). cDNA was synthesised using High Capacity RNA-to-cDNA kit (ThermoScientific).

Real-time PCR detection for both cell culture samples and zebrafish embryos were done using SYBR Green PCR Master Mix for 40 cycles in a Bio-Rad CFX96 thermal cycler. The threshold cycle (Ct) value for each gene was normalised to the Ct value of a housekeeping gene, *EIF1B* and *rpl13* for cell culture and zebrafish respectively. The relative fold changes were calculated using ΔΔCt method. The primer sequences for target genes are listed in Table S1.

### Soft agar assay

Soft agar assay was performed using CytoSelect 96-Well Cell Transformation Assay (Cell Biolabs) according to the manufacturer’s instructions. The final 0.4% agar layer on which cells were grown in corresponds to a Young’s modulus of <2 kPa.^41^ 7 days after seeding, cell colony formation was examined with EVOS Cell Imaging System (ThermoScientific) under 10X magnification. Cell proliferation was measured with CyQuant NF Cell Proliferation Assay Kit (ThermoScientific) according to manufacturer’s instructions.

### Cell proliferation assay

Cells were plated in 96-well plates at 1,000 cells per well and fluorescent intensity was measured every 24 h post seeding for 7 days according to CyQuant NF Cell Proliferation Assay (ThermoScientific) for adherent cells.

### Western blotting

Cells were seeded in 6-well plates at 300,000 cells per well and cell lysate was harvested 48 h post seeding. Nuclear and cytoplasmic fractions were separated using NE-PER Nuclear and Cytoplasmic Extraction Reagents (ThermoScientific) according to manufacturer’s instructions for adherent cells. Protein concentration of both fractions were measured using Pierce BCA Protein Assay Kit (ThermoScientific). 15 ug of protein were separated by 4-20% Mini Protean TGX SDS-polyacrylamide gels (Bio-Rad) at 120 V for 1 h and transferred onto 0.2 um PVDF membranes (Bio-Rad) at 90 V for 1 h. Membranes were blocked with 3% (W/V) BSA in TBST for 1 h at room temperature (RT) and incubated overnight at 4°C with primary antibodies in 3% (W/V) BSA in TBST. Membranes were washed 3 times for 5 min with TBST and incubated with either anti-mouse or rabbit HRP-conjugated secondary antibodies in 3% (W/V) BSA in TBST for 1 h at RT. Membranes were washed 3 times for 5 min with TBST after secondary antibody incubation and chemiluminescence signal was detected using Clarity Western ECL Substrate (Bio-Rad). All relevant information on antibodies used in this study are listed in Table S2.

### Curve fitting

Time constant (τ) of optoYAP import and export from the nucleus was fitted using the curve fitting function *fit* in MATLAB. Nuclear import rate (τ_a_) was fitted with 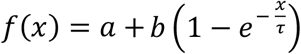 and export rate (τ_r_) was fitted with 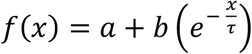.

## Supplementary Methods

**Table S1.**
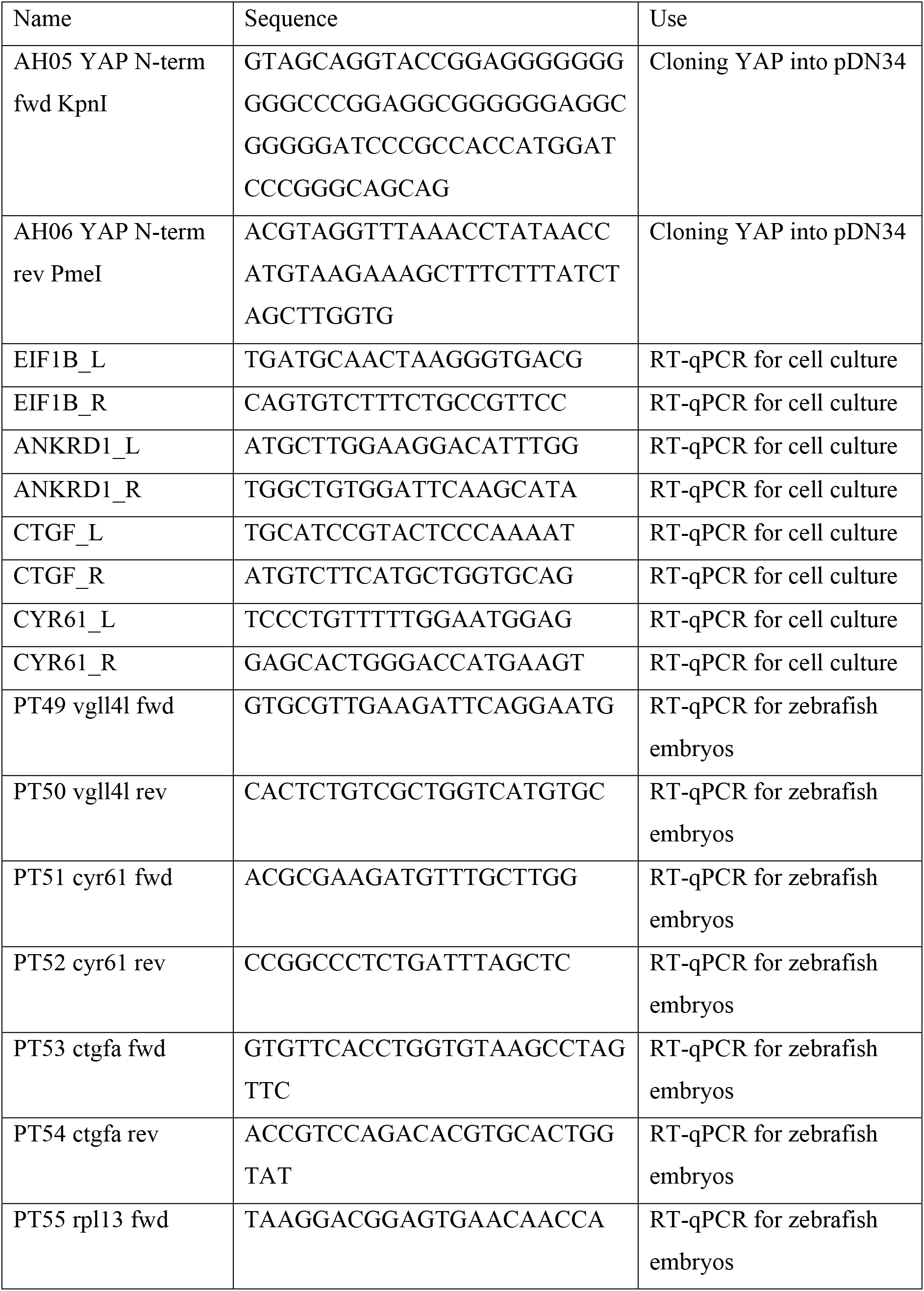
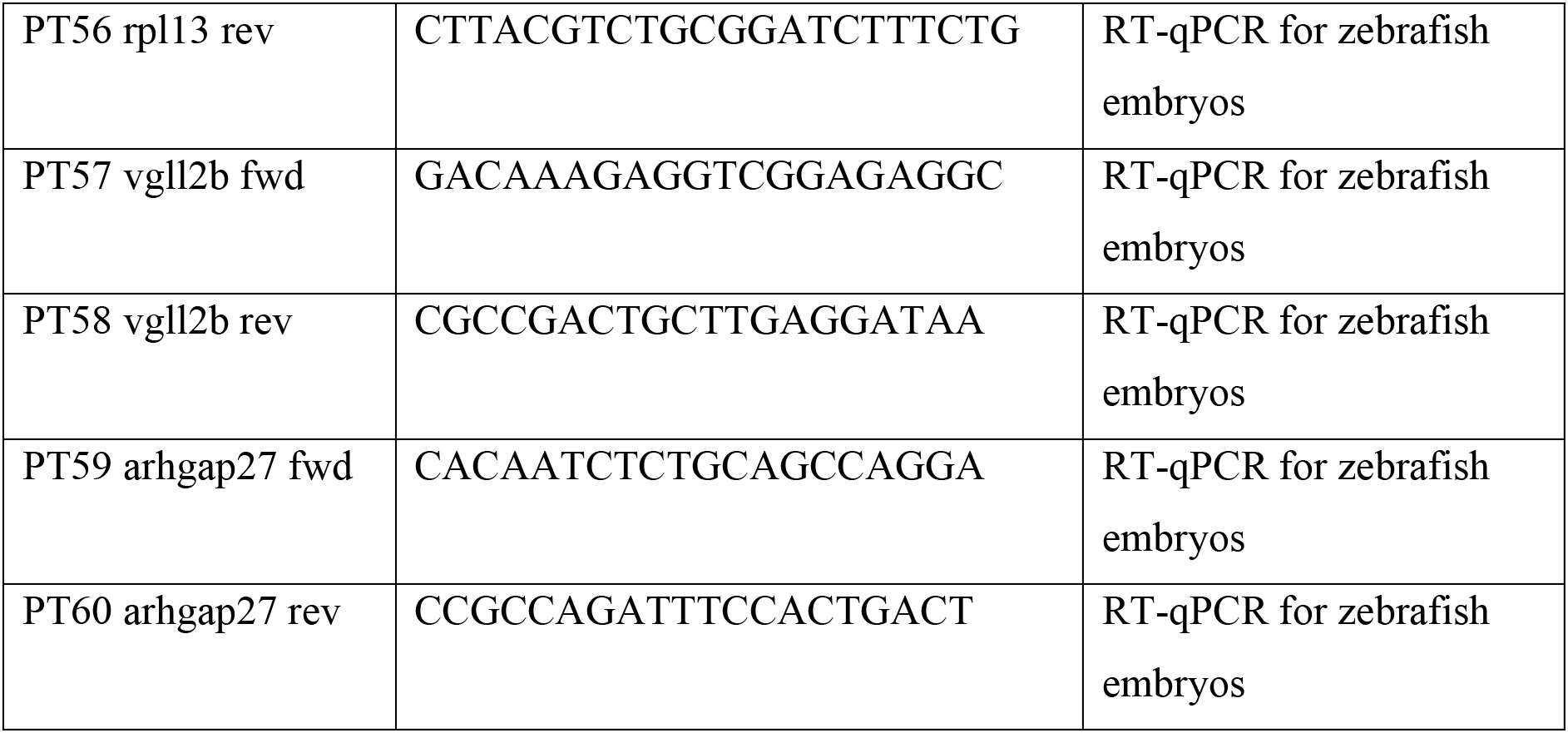
List of primers used.

**Table S2.** List of antibodies used.

| Target protein | Supplier | Catalogue number | Dilution |
| --- | --- | --- | --- |
| YAP (D8H1X) | Cell signalling technology | #14074 | 1:1000 |
| pYAP (S127) | Cell signalling technology | #4911 | 1:1000 |
| GAPDH | Cell signalling technology | #5174 | 1:5000 |
| β-tubulin | Cell signalling technology | #2128 | 1:1000 |
| Lamin B1 | Abcam | ab16048 | 1:5000 |
| Anti-rabbit IgG | Cell signalling technology | #7074 | 1:5000 |
| Anti-mouse IgG | Cell signalling technology | #7076 | 1:5000 |

